# Insect threats and conservation through the lens of global experts

**DOI:** 10.1101/2020.08.28.271494

**Authors:** Marija Miličić, Snežana Popov, Vasco Veiga Branco, Pedro Cardoso

## Abstract

While many recent studies have focused on global insect population trends, all are limited either in space or taxonomic scope. Since global monitoring programs for insects are not implemented, biased data are therefore the norm. However, expert opinion is both valuable and widely available, and should be fully exploited when hard data are not available. Our aim is to use global expert opinion to provide insights on the root causes of potential insect declines worldwide, as well as on effective conservation strategies that could mitigate insect biodiversity loss. We obtained 753 responses from 413 respondents with a wide variety of expertise. The most relevant threats identified through the survey were agriculture and climate change, followed by pollution, while land management and land protection were recognized as the most significant conservation measures. Nevertheless, there were differences across regions and insect groups, reflecting the variability within the most diverse class of living organisms on our planet. Lack of answers for certain biogeographic regions or taxa also reflects the need for research, particularly in less investigated settings. Our results provide a first step towards understanding global threats and conservation measures for insects.

## Introduction

Insects play a key role in providing numerous irreplaceable services, many of which are critical to human survival and wellbeing. Yet, most insects are non-charismatic at best and perceived as pests at worst. As such, they receive little attention, attracting few resources for monitoring and conservation (Krause & Robinson, 2017). Likewise, recent studies on animal biodiversity confirm an under-representation of insects in the published literature (Titley, Snaddon, & Turner, 2017). Nevertheless, declines have not only been observed in relation to insect diversity and abundance (Biesmeijer et al., 2006; Shortall et al., 2009), but also regarding total insect biomass. For example, one pioneer study in Germany showed a dramatic 75% decline in total flying insect biomass, regardless of habitat type (Hallmann et al., 2017). However, all studies quantifying and assessing long-term global trends in insect abundance or diversity to date (e.g. van Klink et al., 2020) are biased, both spatially and taxonomically, often leading to misinterpretation (Simmons et al., 2019; Didham et al., 2020; Montgomery et al., 2020).

As there is uneven coverage of research across taxa, there is also unequal spatial distribution of studies. The highest density of biodiversity research occurs in temperate climates, markedly in Western Europe (Titley et al., 2017). Bearing in mind that an estimated 85% of insect species are found in the tropics and south temperate regions (Stork, 2018), under-representation of biodiversity and conservation studies in these regions could lead to false notions about the state of insect diversity, threats and their global spatial distribution. Hence, there is an urgent need to know the status of insects across the globe.

Global monitoring programs are not implemented, and unbiased data are not available (Cardoso & Leather, 2019). Thus, we must look for alternative strategies to advance our knowledge on insects, namely expert opinion. It frequently plays a fundamental role in conservation, providing substantive information for decision making processes (Cook, Hockings, & Carter, 2010), horizon scanning (Sutherland, Pullin, Dolman, & Knight, 2004) and risk assessments (Patterson, Meek, Strawson, & Liteplo, 2007). Moreover, International Union for the Conservation of Nature (IUCN) species assessments, particularly regarding insects, almost invariably depend on expert scientific judgments as one of the sources of information that is used to measure our progress towards global biodiversity conservation goals.

Our research focuses on the latest insights from experts studying a wide variety of insects around the world. The purpose of this work is to identify the root causes of potential insect declines across different biogeographical regions and taxa based on expert opinion. In addition, expert opinion is also used to identify effective conservation strategies that could mitigate insect biodiversity loss at a global level.

## Material and methods

Our query had 16 questions in six main sections, pertaining to demography, expertise, ecosystem (dis)services, threats, conservation measures and supporting information (Supplementary Material). The first set of questions (1–4) aimed to provide information about the demographic structure of the respondents (gender, experience, education and work responsibilities). Questions 5-7 were intended to reveal their biogeographic and taxonomic expertise, which was used to sub-set the data. In the case of taxonomic groups, we used order-level. The only exception was for ants, which were analysed as a separate group since we considered them to be a distinct entity. Compared to other Hymenoptera, they provide very different services and suffer from different threats. If the respondents’ expertise covered several biogeographic regions or taxonomic groups, they could either select multiple answers if threats and conservation measures were deemed similar; if answers differed between regions or taxonomic groups, they were encouraged to fill the survey multiple times, one for each region or taxon. Questions 8–9 were related with services and disservices provided by insects. Questions 10–12 focused on the threats to insect populations and on their current trends. Questions 13–14 addressed the current and potential conservation measures focused towards insects, including allocating potential funds for future projects. Finally, questions 15 and 16 were intended to provide references that support the previous answers and leave any additional comments.

We created a Google form query with all questions. To ensure geographical coverage, we contacted around 100 entomological societies globally, and asked them to distribute the query among their members. We also contacted approximately 3000 corresponding authors from papers containing the word “insects”, published in international journals in the last 10 years.

To identify the possible influence of demographic structure of the respondents, biogeographic regions and taxa on the answers provided, we performed PerMANOVA with 99999 permutations within the R package vegan (Oksanen et al., 2019), using all answers as response variables for the three analyses. In cases where respondents answered for multiple biogeographic regions or insect taxa, the answers were analyzed separately within each of the selected regions or orders. To avoid dominance in answers coming from the same respondent, we applied a weighted average, where the weight of each answer was divided by the number of queries that respondent had filled (for multiple regions or taxa). A weighted average was also used to overcome possible unbalances in the number of answers for different regions or taxa. Answers to questions regarding threats and conservation of insects ranging from 1–5 were rescaled from 0–1 (1-not relevant at all, 2-little relevance, 3-somewhat relevant, 4-very relevant, 5-extremely relevant). The relevance score average was bootstrapped (random selection of elements with repetition) to obtain upper and lower confidence limits. The procedure was conducted for answers from all regions and taxa together for a global analysis, and then separately for all eight biogeographic regions and all taxa for which more than 10 answers were available (the remaining taxa were grouped under “Other”).

## Results

We obtained 439 responses. Several doubtful answers were eliminated, which included selection of only “unknown relevance” options in the entire survey, selection of all regions and all orders, or the indication that answers relate to non-insects (spiders). In total, the final data set contained 429 responses from 413 participants. When responses that contained selections of multiple regions or taxa were fragmented, we obtained a total of 753 answers.

### Demographic structure of the respondents

Participants in the survey were predominantly male (298), researchers (359), with more than 10 years of experience (291) and with a PhD (318) (Table S1, Supplementary material). The PerMANOVA analysis showed that in most cases, respondent demography did not influence their answers regarding threats and conservation of insects (Table S2, Supplementary material). However, there was within-group variation in answers based on respondents’ education. To account for this, all consecutive analyses were conducted in two ways: 1) analyzing all answers together, and 2) analyzing answers from respondents holding a PhD degree vs. answers from all respondents with other levels of education. Separate analyses are presented in Supplementary material (Figures S1 to S8).

### Answers per region and taxon

Expertise on the Western Palearctic (Figure 1) and on the orders Coleoptera, Lepidoptera and Diptera were dominant (Figure 2). The PerMANOVA analysis showed that there were statistically significant differences in answers based on both biogeographic region and taxon (Table S3, Supplementary material).

**Figure 1.**
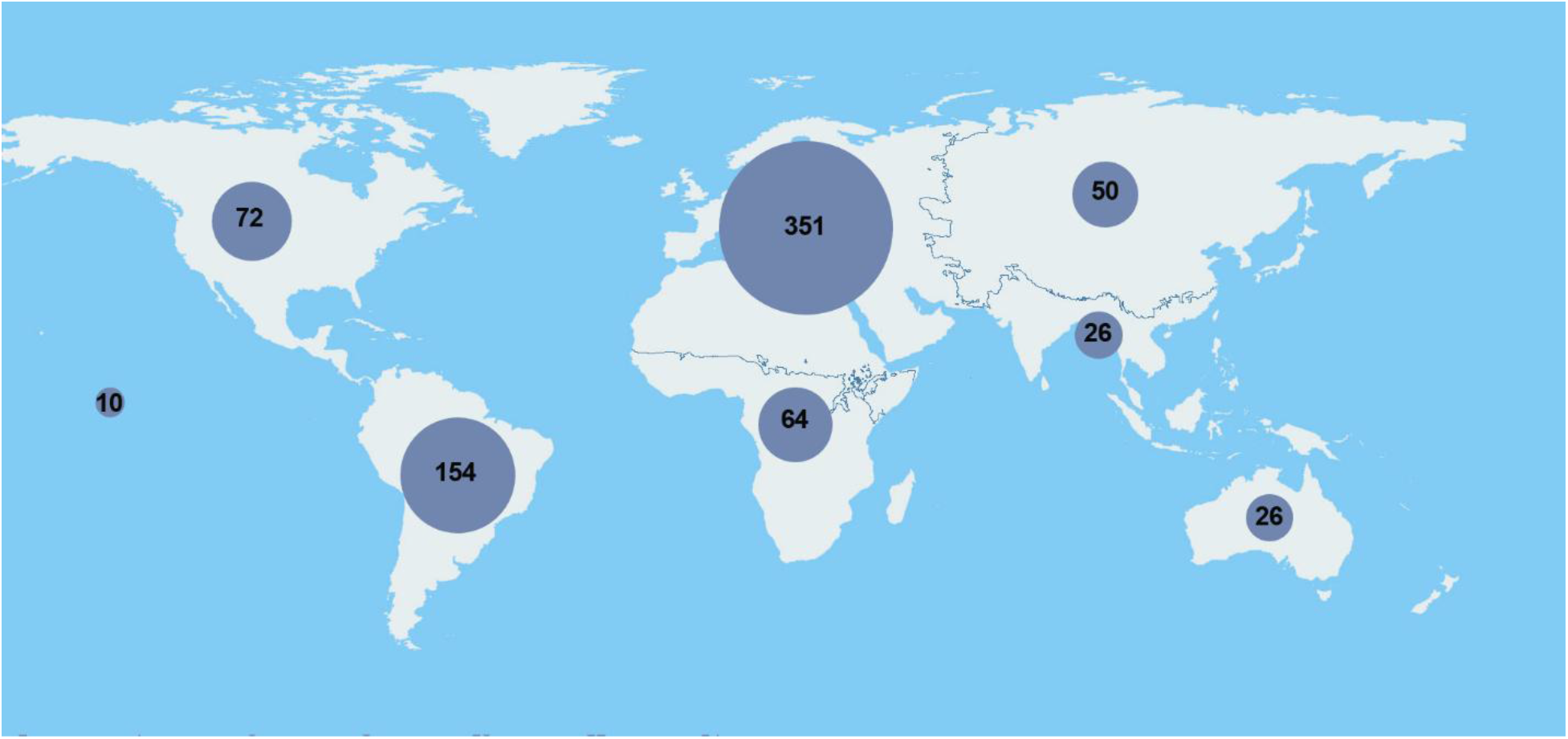
Distribution of answers (with number of respondents) per biogeographical region.

**Figure 2.**
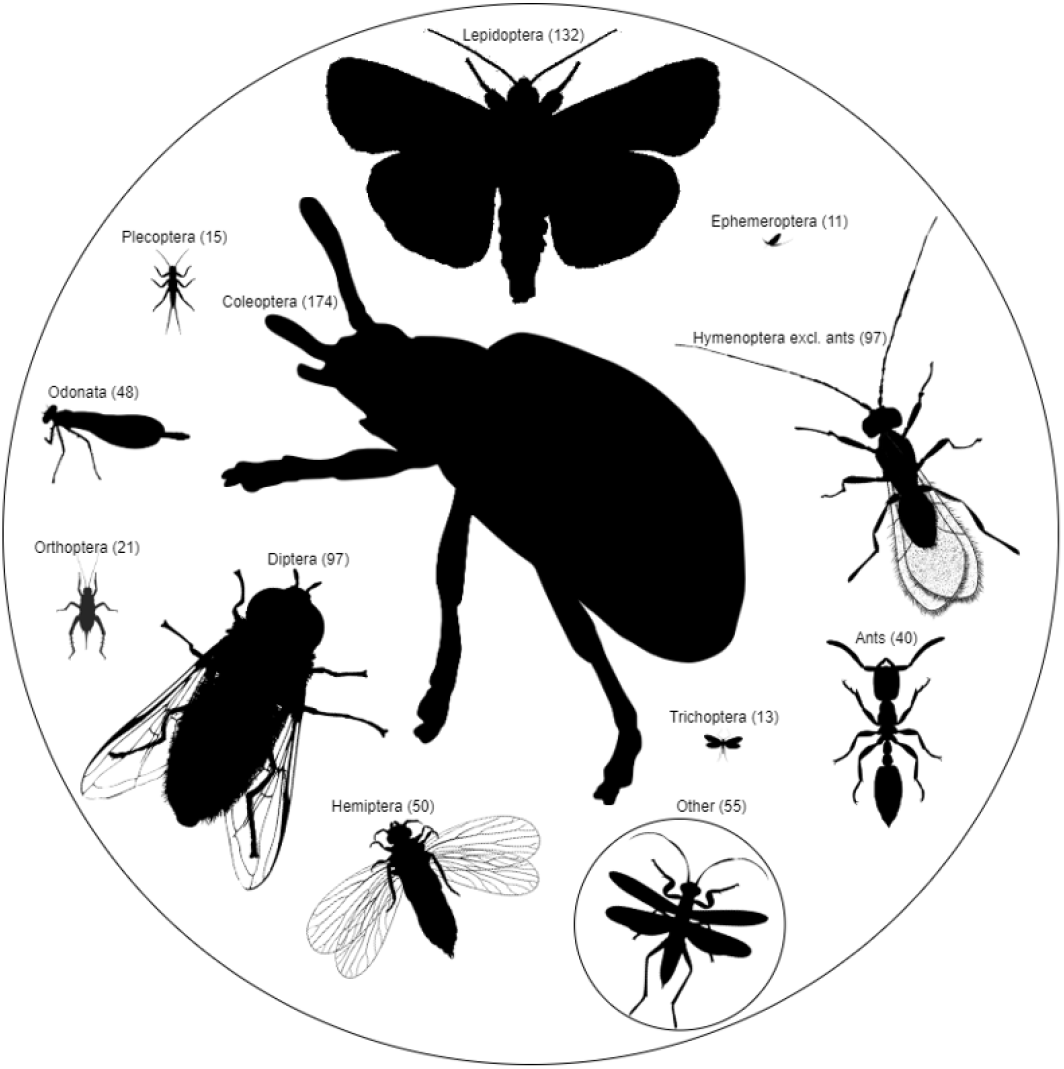
Distribution of answers (with number of respondents) per insect order.

### Insect threats

At a global level, the most relevant threats for insects identified by the survey were agriculture and climate change, followed by pollution, significantly different from the first two (Figure 3). Natural system modifications, invasive species and residential and commercial development were also identified as important, without statistical difference. The Afrotropical, Neotropical, Western Palearctic and Nearctic followed the global pattern regarding the three top-rated threats. In the Eastern Palearctic, Indomalayan, Australasian and Oceanian regions, residential and commercial development was among the top-rated threats. The most noticeable difference was in Oceania, where co-extinctions were listed among the most significant threats, next to those coinciding with the global pattern (Figure 3).

**Figure3.**
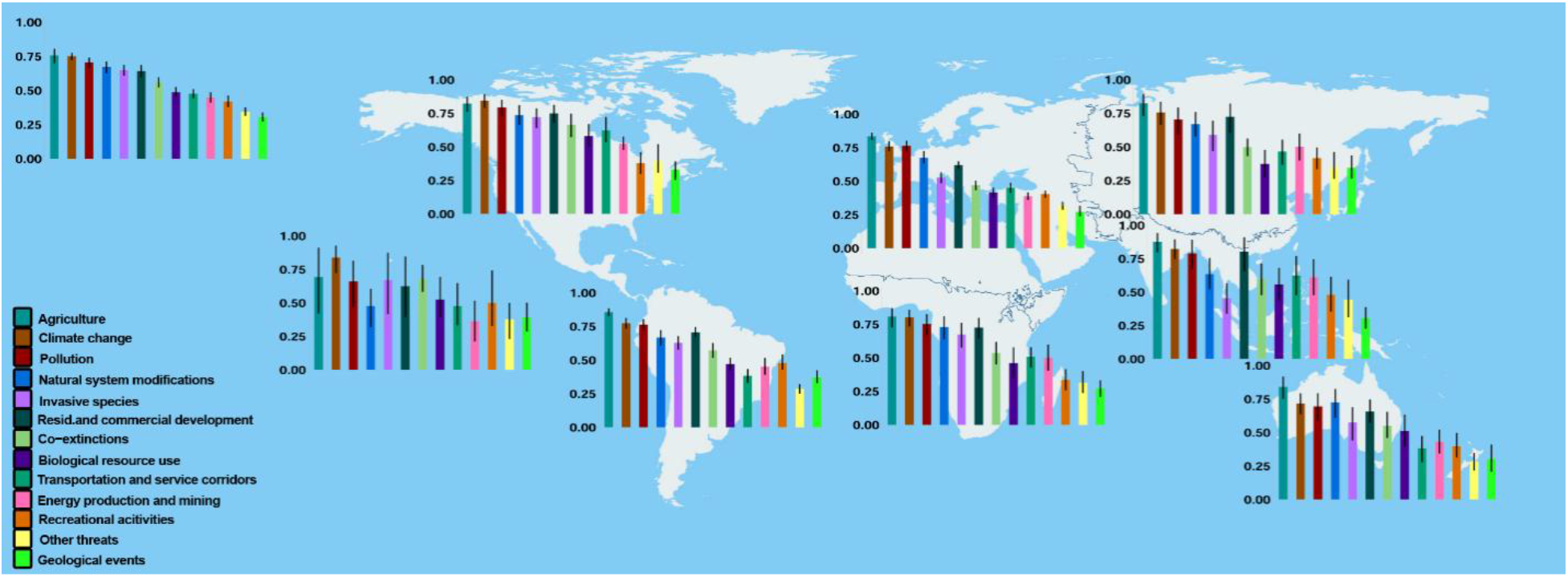
Global (upper left) and biogeographic section-based significance of most relevant treats for insects. Confidence limits were calculated by bootstrap.

In most cases, analyses per insect order revealed that the trends followed the global pattern, with agriculture, climate change and pollution being identified as the most important threats for most orders (Figure 4). For the Ephemeroptera, Plecoptera and Trichoptera (EPT), however, natural system modifications were selected as the most relevant threat. These were also among the most significant responses for Odonata. For ants, in addition to agriculture and natural system modifications, invasive species were among the top-rated threats, not being statistically different from the previous.

**Figure 4.**
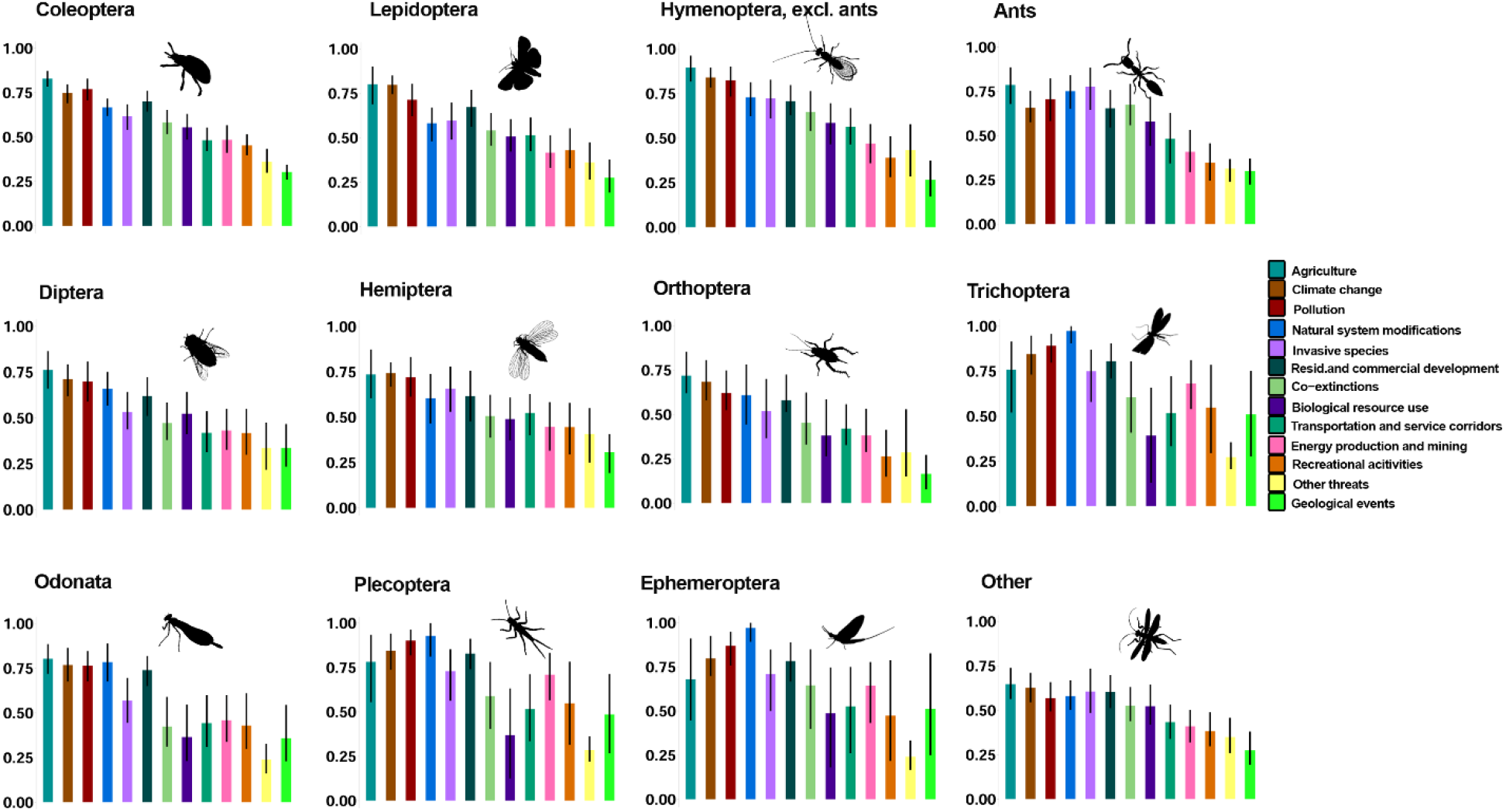
Significance of threatsfor insects by order. Confidence limits were calculated by bootstrap.

There was a perceived decreasing trend in almost all parameters, excluding phylogenetic and functional diversities, for which the trends appeared to be stable or are mostly unknown (Figure 5). It is also worth mentioning that, except in the cases of species richness and abundance, around one third of the respondents marked “unknown trend” for the different measures.

**Figure 5.**
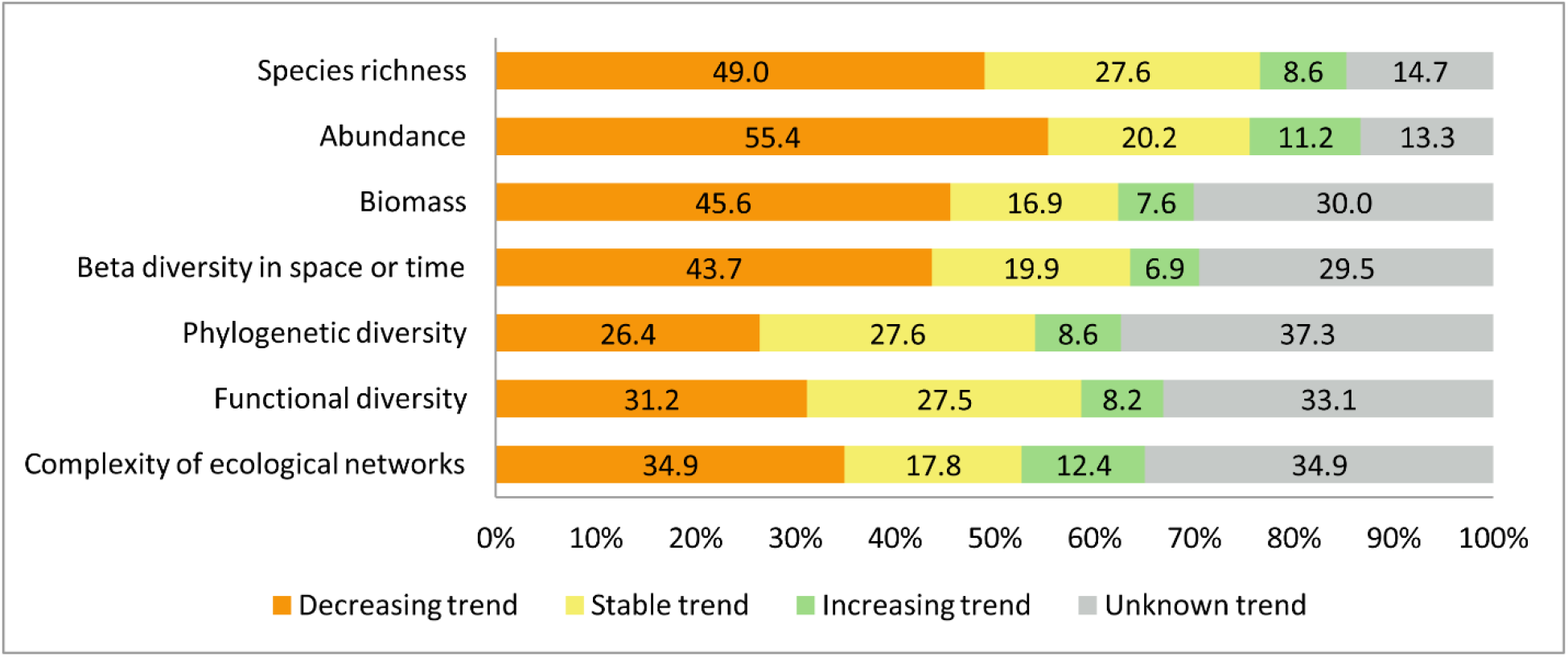
Trends of insect populations. Values are represented in percentages.

### Insect conservation

The conservation measures considered most (and equally) relevant for the global preservation of insects were land management and land protection (Figure 6). In most cases, answers per biogeographic region followed the global trend, with slight but non-significant variations between land protection and land management identified as the most important. However, in the case of the Nearctic, Indomalayan and Eastern Palearctic regions, education was listed among the most relevant. In the Nearctic region, law and policy was also recognized among the most relevant conservation measures (Figure 6).

**Figure 6.**
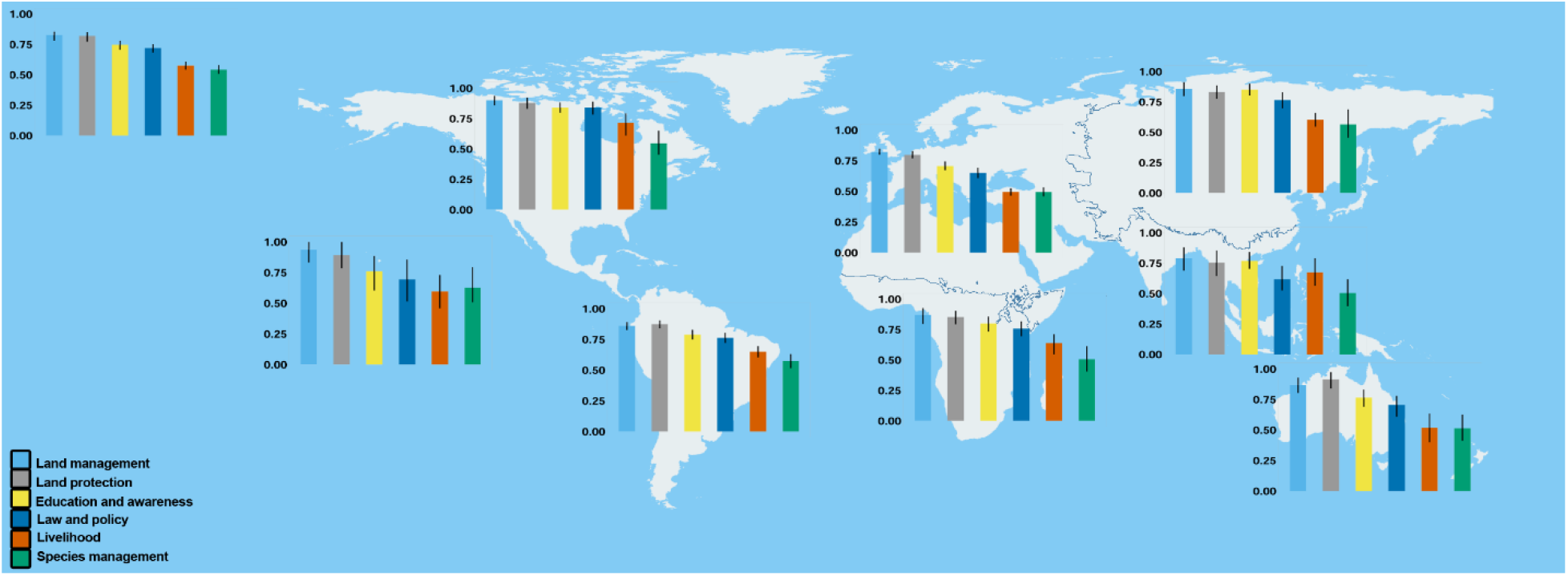
Global (upper left) and biogeographic section-based significance of conservation measures for the preservation of insects. Confidence limits were calculated by bootstrap.

There were some differences in answers regarding the most relevant conservation measures between different insect groups (Figure 7). In Hymenoptera, Hemiptera and Diptera, education was identified among the most relevant conservation measures, next to land protection and land management. In Diptera and Hemiptera, law and policy were also marked as being highly important conservation measures.

**Figure 7.**
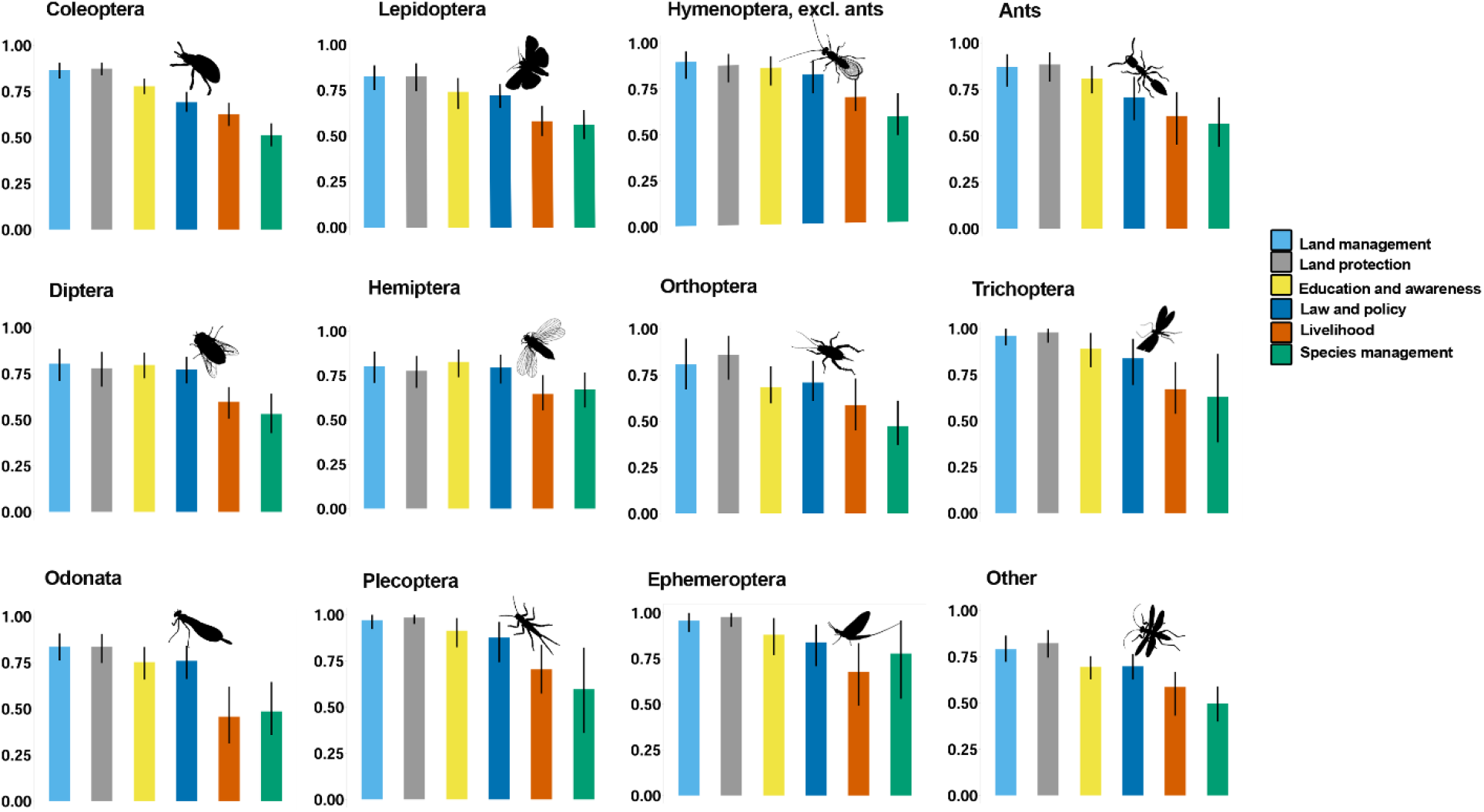
Significance of conservation measures for the preservation of insects by order. Confidence limits were calculated by bootstrap.

Regarding the allocation of the conservation investments, respondents would primarily dedicate funds for research of insect biodiversity, followed by monitoring of biodiversity and acquisition of new protected areas. Management of protected areas, education and management of unprotected areas would follow, while the smallest amount of funds would be allocated to ex-situ conservation of insects (Figure 8).

**Figure 8.**
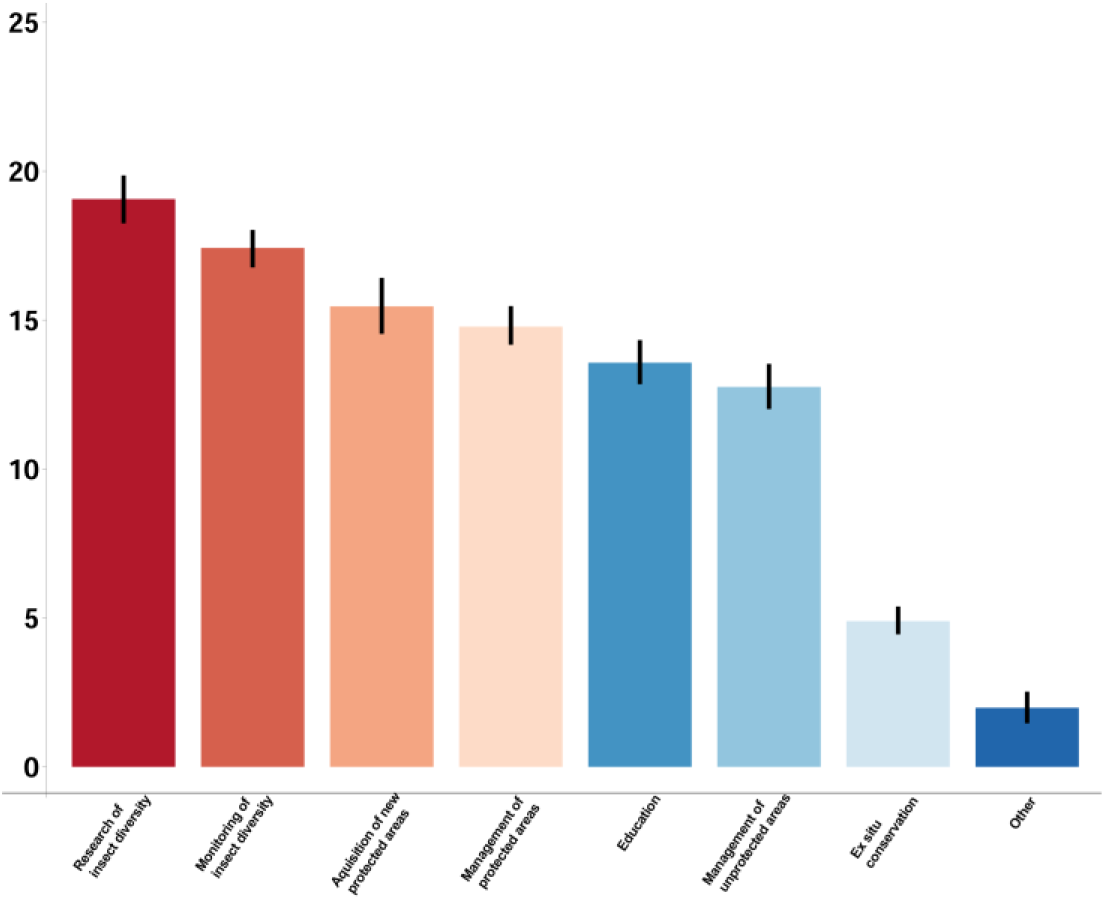
Allocation of funds for conservation investments.

### Insects’ services and disservices

Respondents of the query considered that the most relevant services provided by insects are within provisioning services, namely monitoring of habitat quality and biocontrol (Figure 9). Among regulating services, pollination was depicted as the most significant, while nutrient cycling through saprophagy and coprophagy were the most relevant supporting services. Serving as models for scientific research was the most significant cultural service provided by insects. Pest damage to agriculture and acting as invasive species were selected as main disservices of insects (Figure 10).

**Figure 9.**
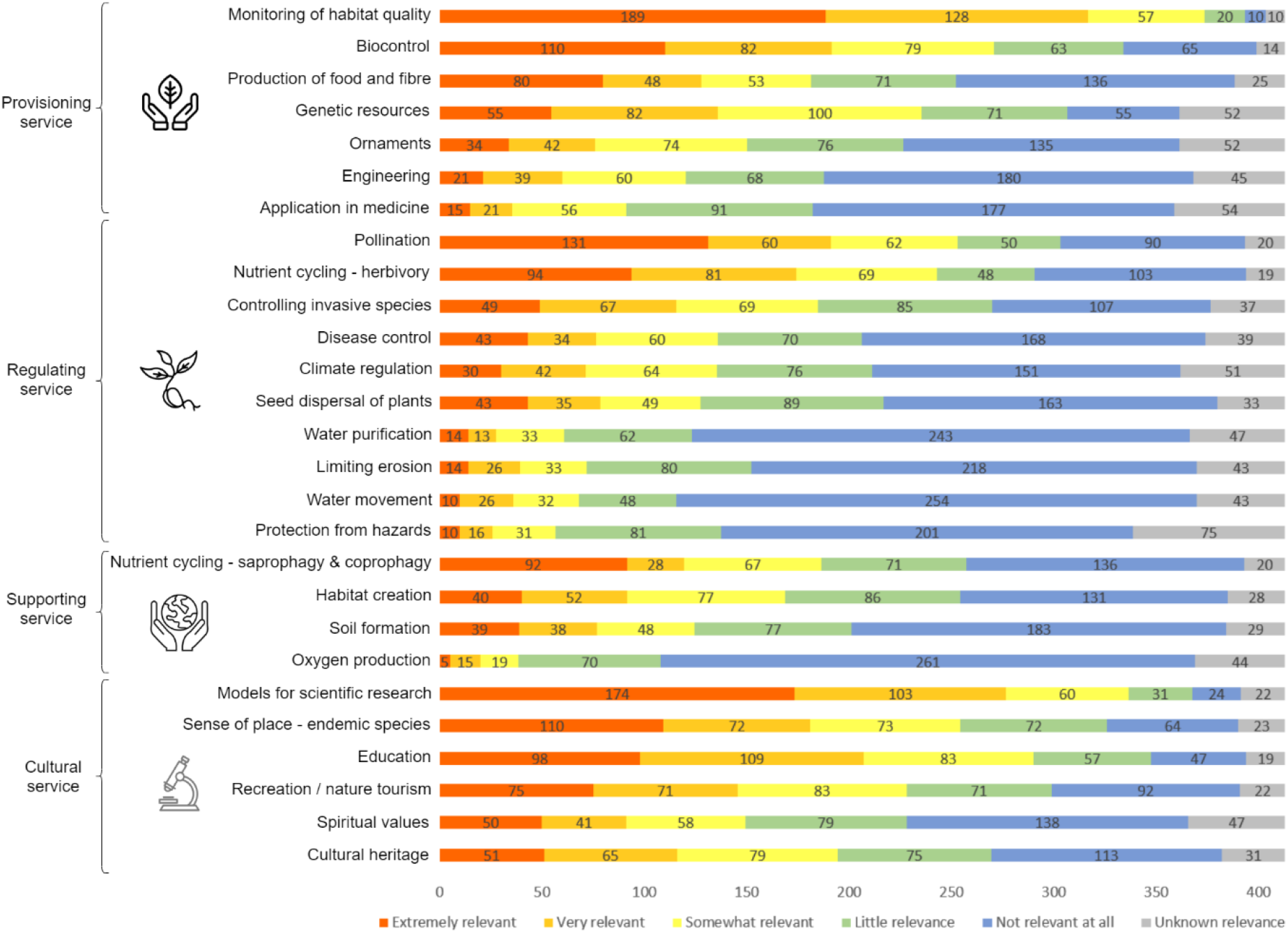
Services provided by insects.

**Figure10.**
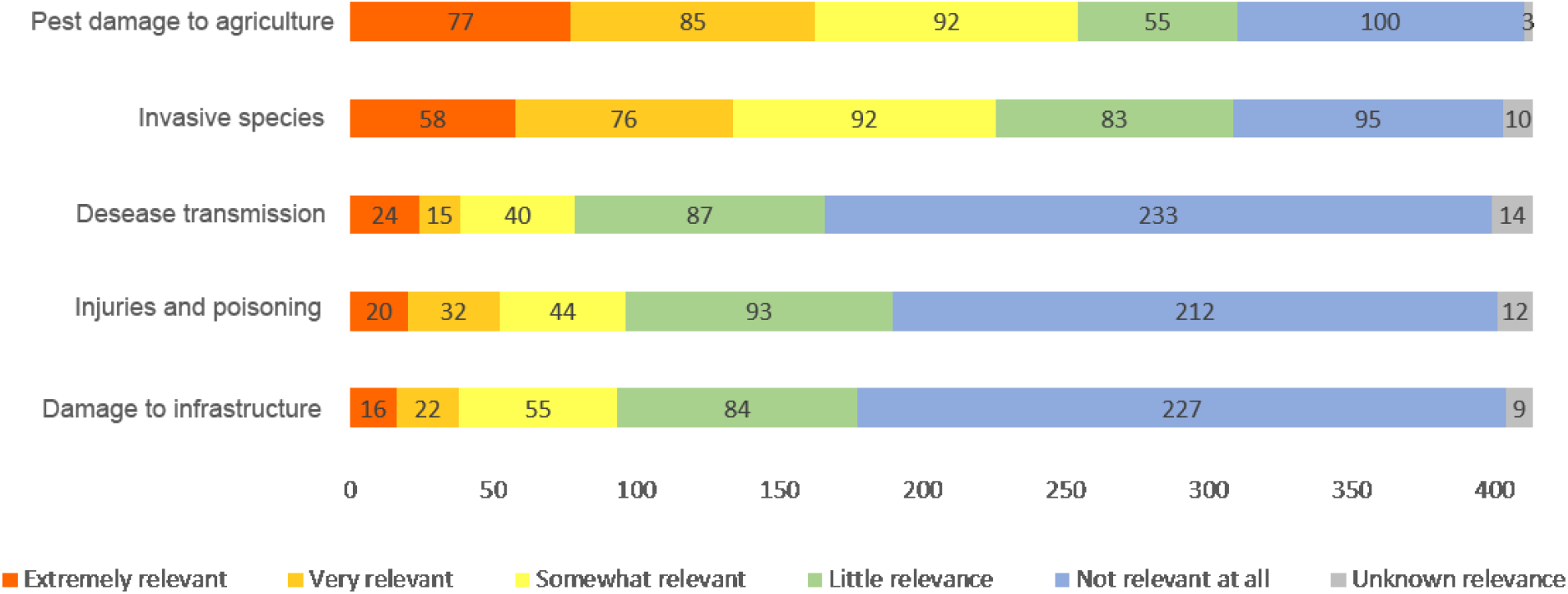
Disservices provided by insects.

## Discussion

Agriculture (including livestock farming) and climate change (including extreme weather events) were consistently identified as the main threats for insects, followed by pollution and natural system modifications. Land management and protection were assigned as the most significant measures that could aid in insect preservation globally. However, there was variation in both the threats and conservation measures identified as the most relevant across different biogeographic regions and taxonomic groups. This suggests that each case must be studied individually when addressing threats and designing conservation actions, depending on the target area or taxonomic group of species.

### Threats

When agriculture and insects are put in the same sentence, one would mainly think of insects as pests, causing economic damage. However, different agricultural activities have proven to be a serious threat not only for insects (Showket, Mir, Parry, Yaqob, & Nisar, 2017), but also to biodiversity worldwide. According to the ESA (Earth Space Agency), about 37% of the entire Earth’s land is used for agricultural purposes; roughly 11% of which used for growing crops, and the remainder for pasture. This type of land transformation leads to habitat loss and biotic homogenization (Olden & Rooney, 2006), both of which subsequently drive the loss of species and facilitate the process of ecological drift, i.e. the replacement of specialists by generalists in species assemblages, as shown by Polus, Vandewoestijne, Choutt, & Baguette (2007) with butterflies in Belgium. Likewise, biotic homogenization through the spread of wood plantations has proven to have negative effects on both taxonomic and functional diversity of Odonata studied in the Neotropics (Dalzochio, Périco, Renner, & Sahlén, 2018) and in the Indo-Malayan region (Dolný, Harabiš, Bárta, Lhota, & Drozd, 2012), which is in accordance with our results. Grazing, as a dominant form of agricultural activity (based on the percentages mentioned above), influences insects in several ways. Primarily, it causes alterations to the plant communities in a system (Börschig, Klein, von Wehrden, & Krauss, 2013), thus causing a cascade effect on many insect populations. It also modifies physical properties of soil and induces changes in microclimate. These effects have proven to promote destruction of nesting sites and removal of potential insect foraging and/or sheltering habitats, as shown for grasshoppers (Orthoptera) (Black, Shepherd, & Vaughan, 2011). Studies in the Neotropical region have highlighted the effect of cattle ranching on the homogenization of the landscape (Kappelle & Brown, 2001), which created more severe environmental conditions through opening of drier open areas. Nichols et al. (2007) relate these conditions to the modified assembly structure of dung beetles (Coleoptera), while Ma et al. (2017) associate the decrease in abundances of two Orthopteran and Hemipteran families in Eastern Palearctic with overgrazing. Arid ecosystems, such as those in the Mediterranean zone and in the Afrotropical region are facing the risk of desertification due to overgrazing, driving semi-arid ecosystems towards desert-like landscapes (Louis, 2016).

Our results on climate change confirm the global concern about the possible consequences of global warming, not only in the scientific literature, but also in mass media (Legagneux et al., 2018). Due to the popularity of the topic, it is not surprising that much research is focused on studying the effects of climate change on living organisms. The most significant effect of climate change on species is the loss of suitable conditions for their survival in a particular ecosystem. We expect that global warming will force species distributions towards “colder” areas at higher elevations and latitudes. Range shifts are documented, for example, in two North American bark beetles (Williams & Liebhold, 2002). Besides range shifts, climate change can cause insects to alter their phenology in order to adapt (Forrest, 2016), either going through evolutionary shifts (Kellermann & van Heerwaarden, 2019) or becoming extinct. However, the biggest concern is that current climate change impacts on biodiversity are several magnitudes lower than those predictable in the future. Although not specifically focused on insects, the work of Kingsford & Watson (2011) and Duffy (2011) address the question of climate change effects in the Oceania region. Duffy (2011) points out the possibility that, due to the all-inclusive consequences of climate change, conservation of biodiversity could be a low priority unless directly related to human needs.

Considering the variety of contaminants in the environment that can threaten insects, it is not surprising that pollution was rated as one of the most serious threats among the respondents. However, certain groups of insects might be more prone to specific types of pollution. For example, a significant number of studies have addressed the effect of light pollution on different beetle species. Not only does it influence nocturnal insects such as fireflies (Lampyridae) by affecting their flash signals (Owens, Meyer-Rochow, & Yang, 2018) and intra- and inter-specific interactions (Firebaugh & Haynes, 2019), but it is also related to the reduced occurrence of fireflies in the Neotropics (Hagen, Santos, Schlindwein, & Viviani, 2015). Besides Coleoptera, pollution was depicted as the second most relevant threat for EPT as well. Considering their dependence on water – at least during part of their life cycle – different water effluents possess a severe threat for these communities, as shown in the study of Chi, Hu, Zheng, & Dong (2017), who found the EPT to be particularly sensitive to the effect of arsenic pollution.

Natural system modifications were the fourth most relevant threat overall; this was second most relevant for Odonata and the most relevant for the EPT species. These four taxa are driving the overall pattern. As previously mentioned, these insect groups are dependent on water bodies, and are proven to be sensitive to human pressure. Thus, it comes as no surprise that water management and building of dams negatively influences these species. Bredenhand & Samways (2009) found that among all analysed macro-invertebrates in Eerste River, South Africa, the EPT community was the most affected by the existence of a dam, as diversity and abundance of the community dropped to almost zero as a result of the impoundment.

In some biogeographic regions or insect groups, certain threats show a contrasting pattern with the global distribution of answers. For example, in the Eastern Palearctic and Indomalayan regions, residential and commercial development was marked as a very significant threat, not significantly lower than the top rated. In the Eastern Palearctic, remarkable economic growth in some regions such as China and Japan, goes hand in hand with infrastructure development, which reflects negatively on biodiversity (Squires, 2014). Investigation of bees in four megacities in Southeast Asia showed that the mean species richness and abundance were significantly higher in the peripheral suburban areas than central business districts (Sing et al., 2016), indicating the sensitivity of this insect group towards urbanization, as also reflected in our results. As for the Indomalayan region, Wattanachaiyingcharoen, Nak-eiam, Phanmuangma, Booninkiaew, & Nimlob (2016) highlighted that a low standard of management in the tourist industry in Thailand may cause an accelerated reduction of the firefly fauna in this region. Furthermore, due to recent unplanned developmental activities, some areas in the Indomalayan region have displayed signs of rapid habitat destruction, negatively affecting biodiversity (Bhattacharya, 2019), which might explain why respondents from this region gave increased weight to this threat. Co-extinctions were depicted as a very important threat in Oceania, not significantly lower than climate change and agriculture. This might be due to the insular nature of this region, which supports high levels of localized endemism (Kier et al., 2009). Thus, extinction of host animals or plants for some insect groups would promote the extinction of dependent insects as well. Oceania was also the region where invasive species were given higher weight. Islands are particularly prone to the effects of invasions due to their spatial and temporal isolation (Borges, Rigal, Ros-Prieto, & Cardoso, 2020), with native species often incapable of adapting to new competitor, predator, or parasite species. Invasive species were also depicted as being a relevant threat for ants. The impact of invasive species on indigenous ants in Indonesia was previously documented by Bos, Tylianakis, Steffan-Dewenter, & Tscharntke (2008). A particularly aggressive ant species, the Argentine ant (*Linephitema humile*) is known to cause major disruptions in native fauna, including outcompeting native ant species (Rowles & O’Dowd, 2007). However, it is worth noting that Oceania was the region with the lowest number of respondents (10). Confidence intervals in this case were proportionally high, so this result should be taken with caution.

Due to the extent and severity of threats that insects are facing worldwide (Cardoso et al., 2020; Wagner, 2020), it was somehow expected that respondents would identify a mainly decreasing trend in parameters reflecting the status of insect populations. However, it is noteworthy that in almost one third of the answers, respondents selected unknown trends, which indicates that, even with the recent rise in the number of papers tackling insect declines, the most diverse class of organisms on the planet is still severely understudied, hence a nuanced view of insect declines or increases according to taxon and region is needed (van Klink et al., 2020). Functional and phylogenetic diversity had more neutral answers, which might reflect the fact that unique functions or branches in the tree of life are not necessarily lost at the same pace as other metrics, not necessarily that they are not lost at all.

### Conservation

Land management and land protection were recognised as the most relevant conservation measures for the global preservation of insects across regions and taxa. Many regions worldwide have been lagging in the protection of relevant areas for insect conservation (Taylor et al., 2018). Taking a landscape perspective to protection and management of insect biotopes is in fact one of the most effective ways to protect countless taxa, their unique ways of life, evolutionary history and complex networks (Samways et al., 2020). Recent findings show that area size, habitat quality and/or suitability are key points when designating zones that are important for conservation of certain insect species (Janković, Miličić, Ačanski, & Vujić, 2020). Increasing the area and connectivity of protected zones promotes richer insect communities, particularly regarding sensitive orders with specific habitat requirements, such as the EPT (South, DeWalt, & Cao, 2019). Likewise, ant species can be protected most easily by protecting the biotopes in which they occur (Mabelis, 2007). Along with ample land protected, it seems that conservation of insect biodiversity also largely depends on managing sufficient and suitable habitats. For instance, land clearing and removal of over-mature and veteran trees have been identified as threats to some of the orders with many species such as Hymenoptera, Coleoptera, Lepidoptera, and Diptera (Sands, 2018). Therefore, land management practises that avoid harvesting large old growth trees would be of conservation importance for many insect species worldwide (Jörg et al., 2016). However, if over-mature trees are present in urban parks, they could be considered as a hazard for public safety and thus get cut by management authorities (Carpaneto, Mazziotta, Coletti, Luiselli, & Audisio, 2010). This is precisely why conservation of insects is not restricted to protected area managers only – tackling this complex challenge requires involvement and collaboration of diverse stakeholder groups, including the general public.

One way of gaining an appreciation of insect diversity is to actively involve citizens in scientific projects. Citizen science projects could greatly increase community-based conservation outcomes, providing data with clear conservation relevance (Oberhauser & Prysby, 2008), as well as favouring connection between citizens and scientists. Acquisition of scientific knowledge through environment-based education programs can be especially powerful in increasing biodiversity awareness (Caro, Mulder, & Moore, 2003), which was also recognised by the respondents highlighting education as a conservation measure of a high relevance. This is of particular importance if focused on early childhood experiences, which are one of the drivers of positive or negative human-insect encounters (Nxumalo & Pacini-Ketchabaw, 2017). Interestingly, education is listed as the second most important conservation measure for the Indomalayan and Eastern Palearctic regions, before land protection (albeit not significantly). Insufficient research attention in these regions when it comes to insects leaves a mark on environmental education activities in the region. For example, as noticed by Corlett (2011), information about the Indomalayan fauna is politically and linguistically fragmented among numerous publications. Often it is not easy to obtain information from conservation literature, as it is more focused on vertebrate species while insects are clearly under-represented (Di Marco et al., 2017). Next to land management and land protection, education and education campaigns as a tool for conservation were particularly recognized by those who responded to surveys on the orders Hemiptera and Diptera. Among Hemiptera, species belonging to the families Coccidae, Diaspididae and Pseudococcidae are known as the most serious pests worldwide (Evans & Dooley, 2013).Certain insect groups like pests may be excluded in insect conservation strategies, reflecting both taxonomic and biological ignorance regarding the importance of these insects. Deeming an insect as a “pest” is simply based on social perception and essentially ignores the key ecological roles played by these species (Delibes-Mateos, Smith, Slobodchikoff, & Swenson, 2011). Similarly, the association of many Diptera species with coprophagy (Khofar, Kurahashi, Zainal, Isa, & Heo, 2019), saprophagy, or disease transmission (Sarwar, 2016) makes their role in nutrient recycling, pollination and biological control often overlooked. It seems that ignorance could be listed as one of the leading threats when speaking about insect species. Thus, dealing with large gaps in taxonomical, biological and ecological knowledge of insects should be of high priority on all educational levels.

According to Samways (2018), currently insect conservation has a foothold in six interconnected themes: philosophy, research, psychology, practice, validation and policy, the latter of which makes the framework for action. The respondents recognized law and policy as a significant conservation measure in all regions and for all taxa. As an example, protection of insect species at risk, as well as conservation of their natural habitats, was recognized as a strong desire of people in the study of Danks, Smith, Foottit, & Adler (2017). Respondents highlighted the importance of law and policy especially regarding Diptera, Hemiptera and Odonata. Worldwide, dragonflies play a central role in the conservation of freshwater habitats, being used as indicators for environmental health and conservation management (e.g. Chovanec & Waringer, 2005). Yet, a recent study (Kalkman et al., 2018) highlighted a strong mismatch between the distributions of species listed as threatened in the IUCN European Red List and species protected by the EU Habitats Directive. The authors call for re-evaluation of the list of species included in the Directive so it can include Mediterranean threatened species that are neglected under the current policy. The same could be said of other insects and arthropods, with the big, beautiful and common species being privileged in their protection at the European level (Cardoso, Erwin, Borges, & New, 2011) at the expense of the small and “ugly” ones like Diptera and Hemiptera.

According to the respondents, the least relevant among all conservation measures was species management. A common challenge in insect species conservation and management is how to manage species of which our knowledge is limited, particularly in relation to life-cycle interventions, which is a pre-requisite for the successful implementation of management actions. Moreover, handling the enormous richness of insect orders makes it difficult to manage species individually. Nevertheless, species management may be a tool to be explored further in insect conservation.

Regarding the allocation of conservation investments, the respondents indicated that they would primarily fund research and monitoring of insect biodiversity. Incorporating insects into protected area monitoring activities is possible only with well-documented species inventories (McGeoch et al., 2011), which brings us back to the recognized gaps in taxonomical, biological and ecological knowledge of insects and the need to build unbiased monitoring programs at a global scale (Cardoso & Leather, 2019). Research studies conducted on a variety of spatial and temporal scales are of crucial importance to gather the data needed to inform insect conservation policy and management and, thus properly address the drivers of insect declines. This is the only way to stop insect declines in view of necessarily limited resources for conservation of countless species, described and undescribed.

### Services and disservices

As insects play a key role in various ecosystem services, further research should also be devoted to service quantification. A recent study shows that the most studied ecosystem services provided by insects are pollination, biological control, food provisioning and recycling organic matter (Noriega et al., 2018). Indeed, the respondents selected biocontrol and monitoring of habitat quality as the most relevant services among provisioning services. Among the regulating and supporting services, pollination and nutrient cycling through saprophagy and coprophagy were selected as the most significant. Interestingly, the respondents recognized serving as models for scientific research as the most significant cultural service provided by insects, which is proved by many recent studies worldwide, including evolutionary (Wang et al., 2016), toxicology (Coates et al., 2019), biomonitoring (Miguel, Oliveira-Junior, Ligeiro, & Juen, 2017), robotics (Mintchev, de Rivaz, & Floreano, 2017) and many other studies in different areas. As for disservices, pest damage to agriculture and acting as invasive species were listed as the most significant. The flows of services and disservices, at least in agricultural ecosystems, directly depend on how ecosystems are managed at different landscape scales (Zhang, Ricketts, Kremen, Carney, & Swinton, 2007). Indeed, this should be definitely taken into account when planning policy-relevant entomological research that can influence agricultural practices. The identification of potentially high-risk invasive pest species is the first step in prioritizing species as candidates for further research (Worner & Gevrey, 2006).

### Conclusion

Our study shows that experts consider agriculture, climate change and pollution as the three main drivers for insect declines across different biogeographic regions and orders. Land protection and land management are identified as the most prominent conservation measures that might aid in the preservation of insects and mitigation of negative consequences worldwide. A decreasing trend in parameters reflecting the status of insect populations, as well as the extreme significance of insects, as reflected in the numerous ecosystem services they provide, calls for global monitoring programs and a specialized approach in conservation for each particular case.

## Acknowledgements

We thank Marina Szulc for the help and advice regarding data analysis and visualization. We give attribution to George Starr (Ephemeroptera, link to license https://creativecommons.org/licenses/by-nc-sa/3.0/), Maxime Dahirel (Odonata, https://creativecommons.org/licenses/by/3.0/), Melissa Broussard (Orthoptera, https://creativecommons.org/licenses/by/3.0/), Nico Muñoz (Pleoptera, https://creativecommons.org/licenses/by-nc/3.0/), Didier Descouens (vectorized by T. Michael Keesey) (Trichoptera, https://creativecommons.org/licenses/by-sa/3.0/) for material downloaded from website Phylopic. MM and SP acknowledge financial support of the Ministry of Education, Science and Technological Development of the Republic of Serbia (Grant Nos. 451-03-68/2020-14/ 200358 and 451-03-68/2020-14/ 200125) and H2020 Project ANTARES, grant no. 664387. VVB and PC are supported by Kone Foundation.

## Confilct of interest

The authors declare no conflict of interest.

