## Supplementary material for "Insect threats and conservation through the lens of global experts"

#### Part A - Questionaire

1. What is your gender?
   - Female
   - Male
   - Prefer to self-describe
   - Prefer not to say

2. For how long have you held your insect expertise?
   - Less than 5 years.
   - Between 5 and 10 years.
   - Between 10 and 20 years.
   - Over 20 years.

3. What is your highest level of completed education?
   - Secondary
   - University/College (Bachelor)
   - Masters degree
   - Doctorate
   - Other

4. Indicate the option that best reflects the majority of your responsibilities.
   - Manager / Decision Maker
   - Researcher / Scientist
   - Applied Practitioner / Planner
   - Advocate / Activist
   - Educator
   - Volunteer / Amateur entomologist

5. To what biogeographic area does your study region belong to?
   - Afrotropical
   - Australasian
   - Eastern Palaearctic
   - Indo-Malayan
   - Nearctic
   - Neotropical

- Western Palaearctic
- Oceania

6. Which specific area does your insect expertise cover (please specify continent, country or name of smaller area)? Please answer all following questions in relation to this study area only. If your expertise covers several distinct areas for which the answers would differ, please fill this same questionnaire again, for each area separately.

7. To which insect order your target insect group belongs to? If your expertise covers several distinct orders for which the threats differ, please fill this same questionnaire again, for each order separately. Please, do bear in mind that all your answers should be related to your specific expertise, even if it covers lower taxonomic rank than order.

- Ants
- Archaeognatha
- Blattodea
- Coleoptera
- Dermaptera
- Diptera
- Embioptera
- Ephemeroptera
- Grylloblattodea
- Hemiptera
- Hymenoptera, excluding ants
- Lepidoptera
- Mantodea
- Mantophasmatodea
- Mecoptera
- Megaloptera
- Neuroptera
- Odonata
- Orthoptera
- Phasmatodea
- Phthiraptera
- Plecoptera
- Psocoptera
- Raphidioptera
- Siphonaptera
- Strepsiptera
- Thysanoptera
- Thysanura
- Trichoptera
- Zoraptera
- Other

8. How relevant is the contribution of your target insect group to the following ecosystem services?
    - Provisioning service - Application in medicine
    - Provisioning service – Engineering
    - Provisioning service - Monitoring of habitat quality
    - Provisioning service - Genetic resources
    - Provisioning service – Ornaments
    - Provisioning service – Biocontrol
    - Provisioning service - Production of food and fibre
    - Regulating service - Climate regulation
    - Regulating service - Disease control
    - Regulating service - Limiting erosion
    - Regulating service - Controlling invasive species
    - Regulating service - Nutrient cycling (herbivory)
    - Regulating service - Protection from hazards
    - Regulating service – Pollination
    - Regulating service - Seed dispersal of plants
    - Regulating service - Regulating water movement
    - Regulating service - Water purification
    - Supporting service - Nutrient cycling (saprophagy/coprophagy)
    - Supporting service - Oxygen production
    - Supporting service - Habitat creation
    - Supporting service - Soil formation
    - Cultural service - Cultural heritage
    - Cultural service – Education
    - Cultural service - Models for scientific research
    - Cultural service - Recreation (nature tourism)
    - Cultural service - Sense of place (endemic species)
    - Cultural service - Spiritual values

9. How relevant are the following negative impacts of your target insect group to human health and/or economy?
    - Injuries and poisoning
    - Disease transmission
    - Damage to infrastructure
    - Pest damage to agriculture
    - Invasive species

10. In your research area, how relevant is each of the following as threats to current insect populations?
    - Residential and commercial development
    - Agriculture (e.g. arable farming) / Aquaculture / Livestock farming (e.g. grazing)
    - Energy production and mining (e.g. turbines)

- Transportation and service corridors (e.g. power lines, road lighting)
- Biological resource use (e.g. hunting & collecting, logging & wood harvesting)
- Natural system modifications (e.g. fire or fire suppression, dams and water management/use)
- Invasive alien and other problematic species (e.g. predation by other species, competition with other species, adverse habitat alteration by other species)
- Co-extinctions (of symbionts, hosts or other dependent species)
- Pollution (e.g. pesticides, industrial pollution, domestic and urban waste water, light pollution)
- Climate change and severe weather
- Recreational acitivities
- Geological Events
- Other threats

11. If you considered "other threats" as somewhat relevant or with higher relevance in the previous question, please state these threats here. Are there any other relevant future threats to the insects in your area not stated in the previous question?

12. How is the current trend of your insect group manifested?
    - Species richness
    - Abundance
    - Biomass
    - Beta diversity in space or time
    - Phylogenetic diversity
    - Functional diversity
    - Complexity of ecological networks

13. In your research area, how relevant is each of the following as conservation measures to current insect populations?
    - Land protection
    - Land management
    - Species management (e.g. harvest, trade, ex-situ conservation)
    - Education & awareness
    - Law & policy
    - Livelihood, economic & other incentives

14. Imagine you have to decide how to allocate limited amount of conservation investments to minimize the loss of insect biodiversity. How would you divide those funds? The total must sum to 100%.
    - Research of insect biodiversity
    - Monitoring of insect biodiversity

- Acquisition of new protected areas
- Management of protected areas
- Management of unprotected areas
- Ex-situ conservation
- Education
- Other

15. Please provide some references to published sources that support your answers, if there are some. Both references that you have authored or otherwise are acceptable and, if nothing else, personal communications as well.

16. Please include any additional comments here.

### Part B - Demographic structure and PerMANOVA results

Table S1. Demographic structure of the respondents

| Categories | | Sample size |
|---|---|---|
| **Gender** | | |
| Male | | 298 |
| Female | | 111 |
| Not specified | | 4 |
| | Total | 413 |
| **Education level** | | |
| Secondary education | | 7 |
| Bachelor's degree | | 23 |
| Master's degree | | 65 |
| PhD | | 318 |
| | Total | 413 |
| **Professional designation** | | |
| Researcher | | 359 |
| Volunteer/amateur entomologist | | 22 |
| Manager/decision maker | | 12 |
| Applied practitioner/planner | | 11 |
| Educator | | 9 |
| | Total | 413 |
| **Experience (in years)** | | |
| < 5 | | 30 |
| 5 - 10 | | 92 |
| 10 - 20 | | 122 |
| > 20 | | 169 |
| | Total | 413 |

Table S2. Results of PerMANOVA analysis using demographic structure of respondents in relation to the answers about the threats and conservation measures of insects.

| | Df | Sum Sq | R2 | F | Pr(<F) |
|---|---|---|---|---|---|
| Gender | 3 | 0.0725 | 0.0081 | 1.1626 | 0.2536 |
| Expertise | 1 | 0.0367 | 0.0041 | 1.7667 | 0.1174 |

| | | | | | |
|---|---|---|---|---|---|
| Education | 1 | 0.0653 | 0.0073 | 3.1401 | **0.0150** |
| Work responsibilities | 8 | 0.1506 | 0.0168 | 0.9053 | 0.5285 |
| Residual | 415 | 8.6319 | 0.9637 | | |
| Total | 428 | 8.9571 | 1.0000 | | |

Df - degrees of freedom; Sum Sq - sum of squares; F - F value by permutation, boldface indicates statistical significance with P<0.05. P values are based on 99999 permutations.

Table S3. Results of a PerMANOVA analysis using biogeographic region and insect order in relation to the answers about the threats and conservation measures of insects.

| | Df | Sum of Squares | R2 | F | Pr(<F) |
|---|---|---|---|---|---|
| Region | 8 | 0.7781 | 0.0497 | 4.9583 | **0.0000** |
| Order | 13 | 0.5259 | 0.0336 | 2.0624 | **0.0017** |
| Residual | 731 | 14.3400 | 0.9166 | | |
| Total | 752 | 15.6440 | 1.0000 | | |

Df - degrees of freedom; Sum Sq - sum of squares; F - F value by permutation, boldface indicates statistical significance with P<0.05. P values are based on 99999 permutations.

**Part C - Respondents' answers divided by education**

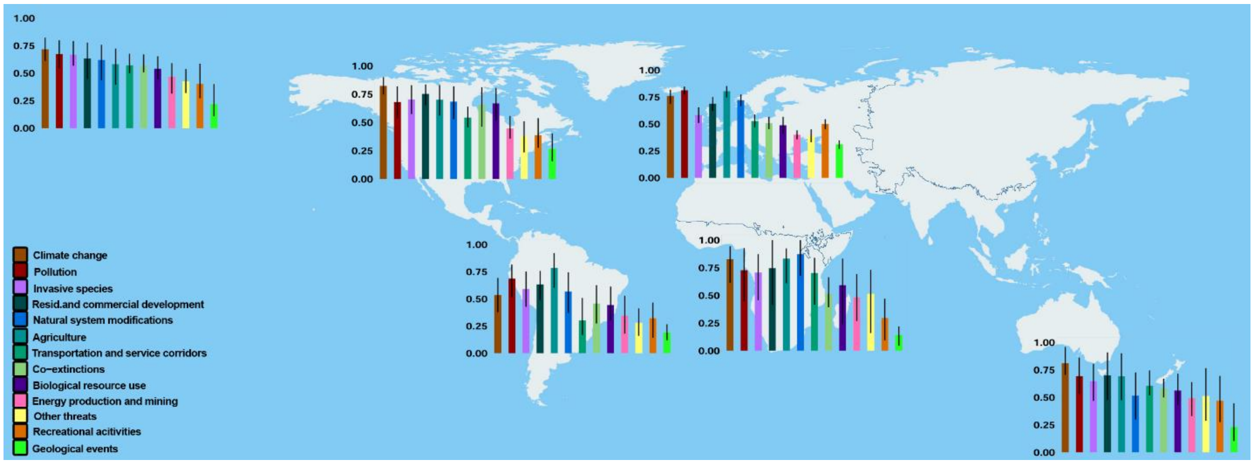


Figure S1. Global (upper left) and biogeographic section-based significance of most relevant treats for insects based on answers of respondents without PhD degree. Confidence limits were calculated by bootstrap. Lower right figure presents the combined answers from all regions for which less than 10 answers were available.

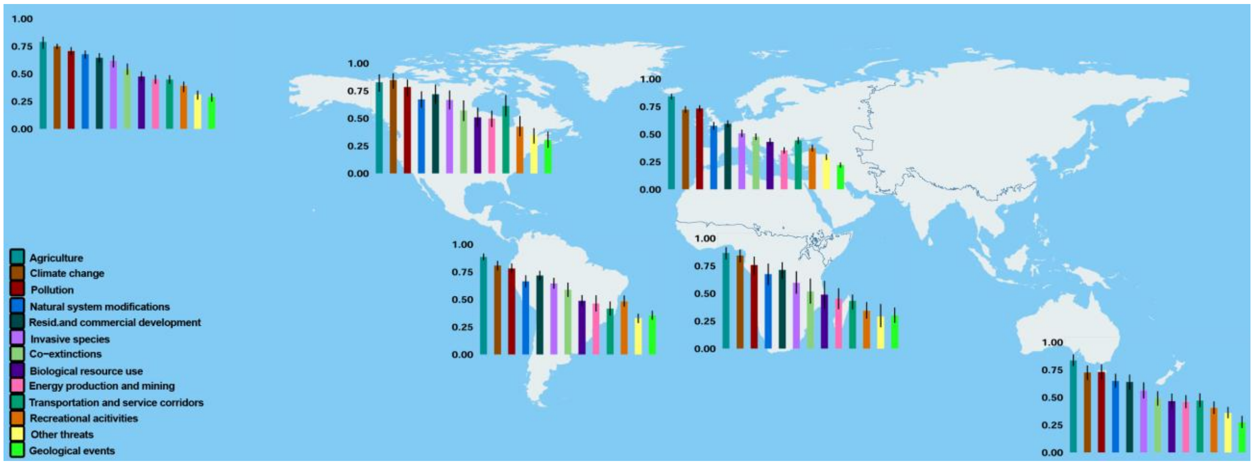


Figure S2. Global (upper left) and biogeographic section-based significance of most relevant treats for insects based on answers of respondents with a PhD degree. Confidence limits were calculated by bootstrap. Lower right figure presents the combined answers from all regions for which less than 10 answers were available.

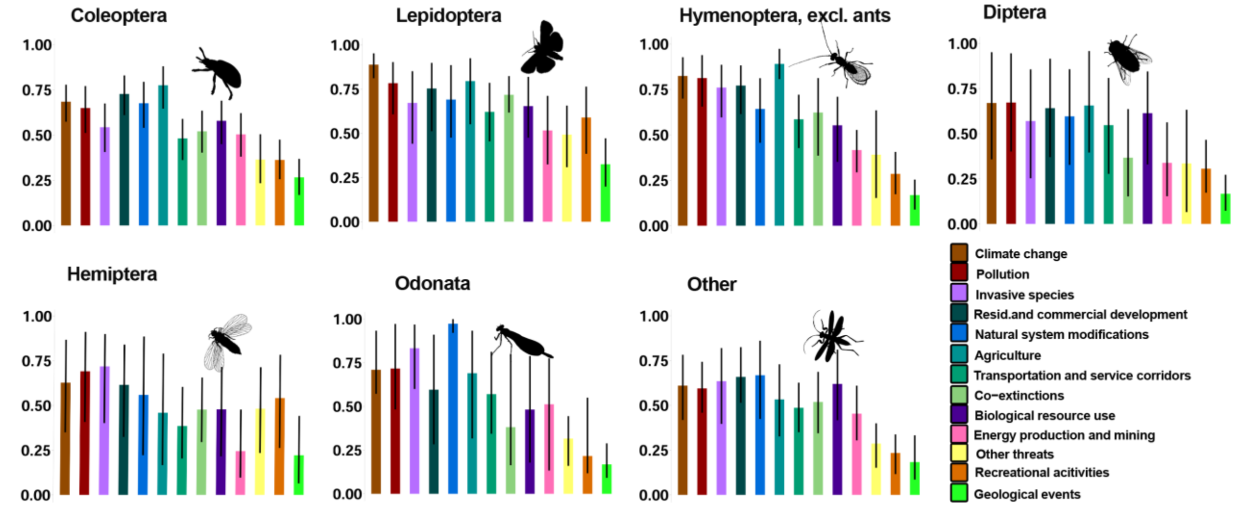


Figure S3. Significance of threats for insects by order based on answers of respondents without PhD degree. Confidence limits were calculated by bootstrap. Figure named “other” presents the combined answers from all orders for which less than 10 answers were available.

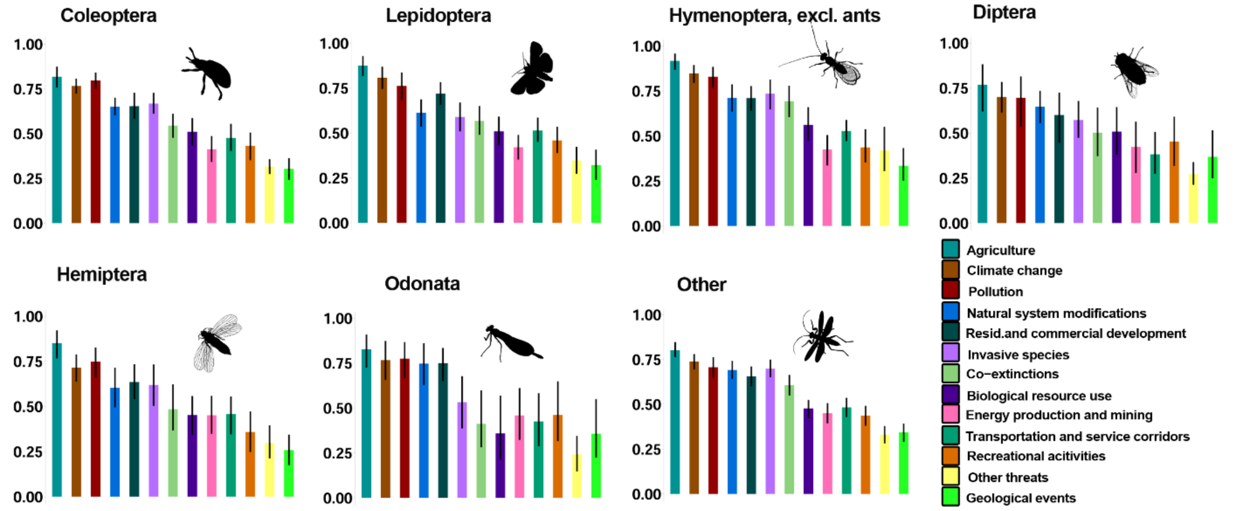


Figure S4. Significance of threats for insects by order based on answers of respondents with a PhD degree. Confidence limits were calculated by bootstrap. Figure named “other” presents the combined answers from all orders for which less than 10 answers were available.

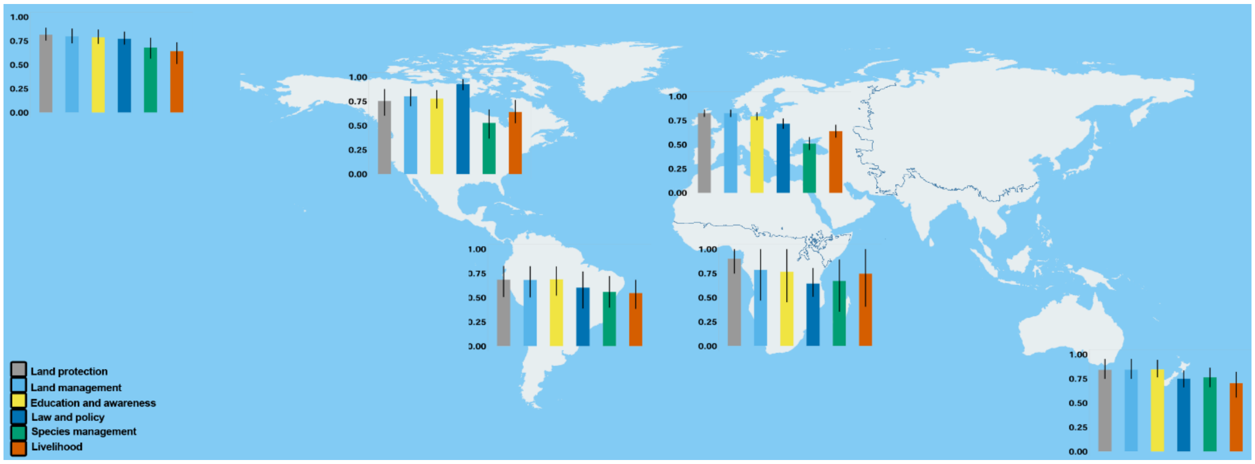


Figure S5. Global (upper left) and biogeographic section-based significance of most relevant conservation measures for insects based on answers of respondents without PhD degree. Confidence limits were calculated by bootstrap. Lower right figure presents the combined answers from all regions for which less than 10 answers were available.

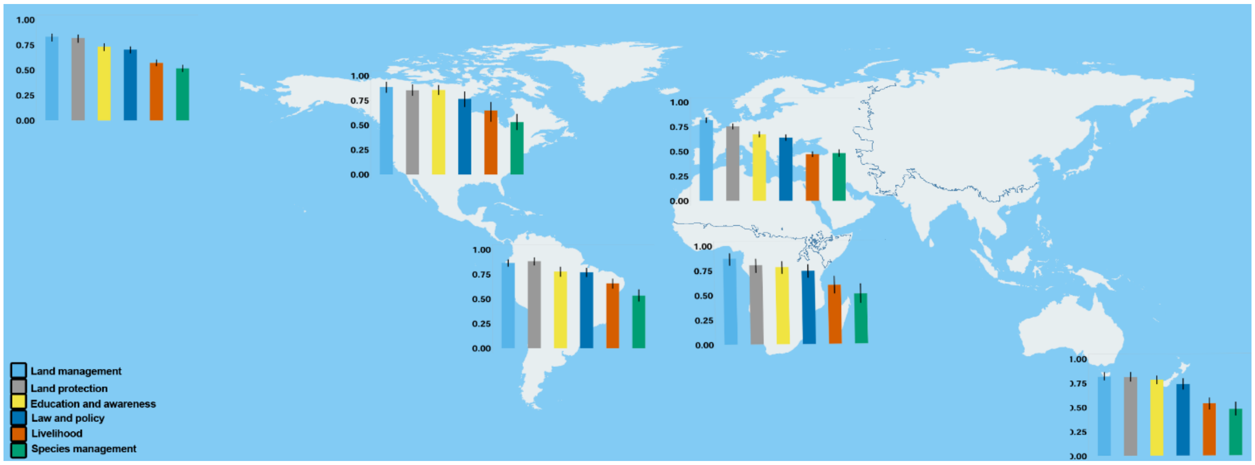


Figure S6. Global (upper left) and biogeographic section-based significance of most relevant conservation measures for insects based on answers of respondents with a PhD degree. Confidence limits were calculated by bootstrap. Lower right figure presents the combined answers from all regions for which less than 10 answers were available.

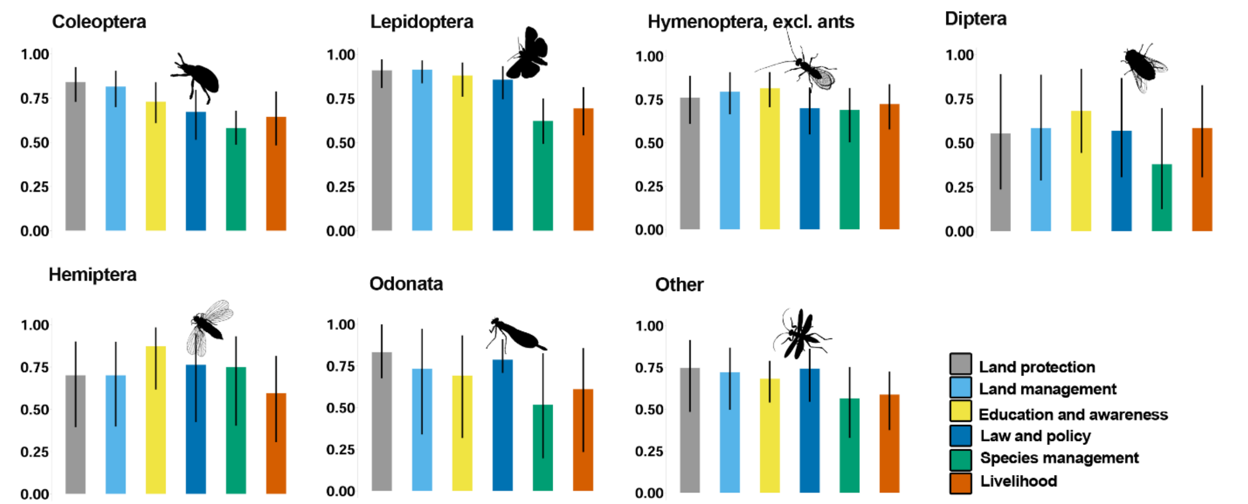


Figure S7. Significance of conservation measures by order for insects based on answers of respondents without PhD degree. Confidence limits were calculated by bootstrap. Figure named “other” presents the combined answers from all orders for which less than 10 answers were available.

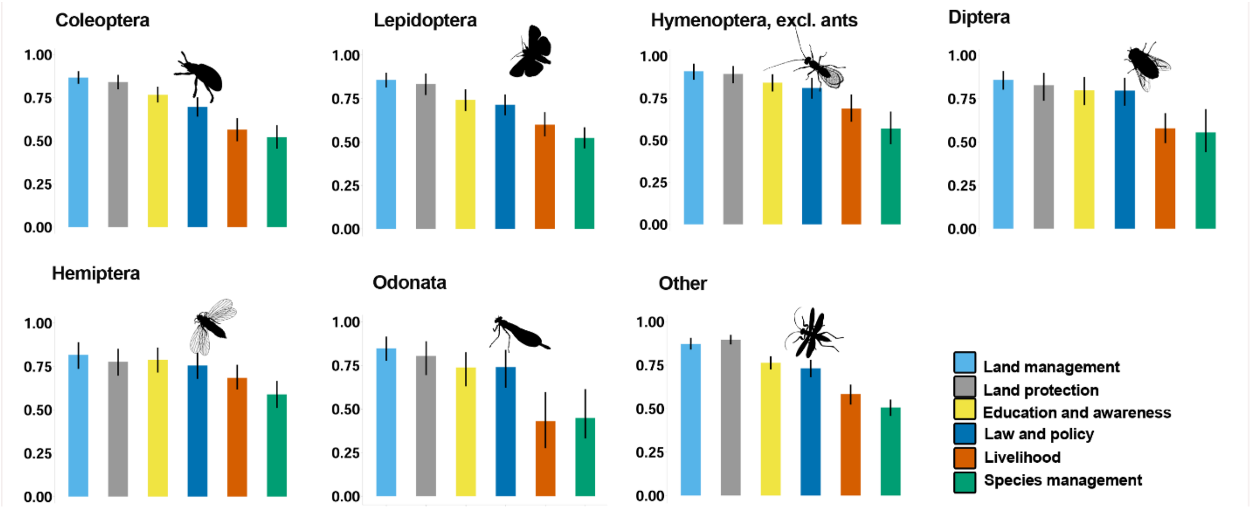


Figure S8. Significance of conservation measures by order for insects based on answers of respondents with a PhD degree. Confidence limits were calculated by bootstrap. Figure named “other” presents the combined answers from all orders for which less than 10 answers were available.
